# Surface-Tethering Enhances Precision in Measuring Diffusion Within 3D Protein Condensates

**DOI:** 10.1101/2025.06.11.659185

**Authors:** Emily R. Sumrall, Guoming Gao, Shelby Stakenas, Nils G. Walter

**Affiliations:** Biophysics Graduate Program, University of Michigan, Ann Arbor, MI 48109, USA; Center for RNA Biomedicine, University of Michigan, Ann Arbor, MI 48109, USA; Division of Biology and Biological Engineering, California Institute of Technology, Pasadena, CA 91125, USA; Department of Chemistry, University of Michigan, Ann Arbor, MI 48109, USA

**Keywords:** biomolecular condensates, membraneless organelles, single-molecule tracking, diffusion, surface passivation, FUS, RNA

## Abstract

Biomolecular condensates, or membraneless organelles, play pivotal roles in cellular organization by compartmentalizing biochemical reactions and regulating diverse processes such as RNA metabolism, signal transduction, and stress response. Super-resolved imaging and single-molecule tracking are essential for probing the internal dynamics of these condensates, yet intrinsic Brownian (thermal capillary wave) motion of the entire condensate in vitro could introduce artifacts into diffusion measurements, confounding the interpretation of molecular mobility. Here, we systematically assess and address this question using both experiments and simulations. We deploy three surface-tethering strategies—using biotinylated DNA, protein, or antibody tethers—to immobilize FUS protein condensates on passivated glass surfaces. We show that tethering effectively suppresses the global translational and rotational Brownian motion of the entire condensate, eliminating inherent measurement artifacts while preserving their spherical appearance and native liquid-like properties. Quantitative analysis reveals that untethered condensates systematically overestimate or underestimate molecular diffusion coefficients and step sizes, particularly for slowly diffusing structured mRNAs, while rapidly diffusing unstructured RNAs are unaffected due to temporal scale separation. Comparative evaluation of tethering strategies demonstrates tunable control over condensate stability and internal dynamics, with implications for optimizing experimental design. Finally, simulations spanning the full physiological parameter space enable us to provide practical guidelines for assessing whether, and to what extent, tethering is beneficial, based on condensate size and the diffusion properties of the biomolecule of interest. Our findings establish surface tethering as a necessary and robust approach for accurate quantification of intra-condensate molecular dynamics, providing a methodological framework for future studies of membraneless organelles.

**TOC Graphic:** 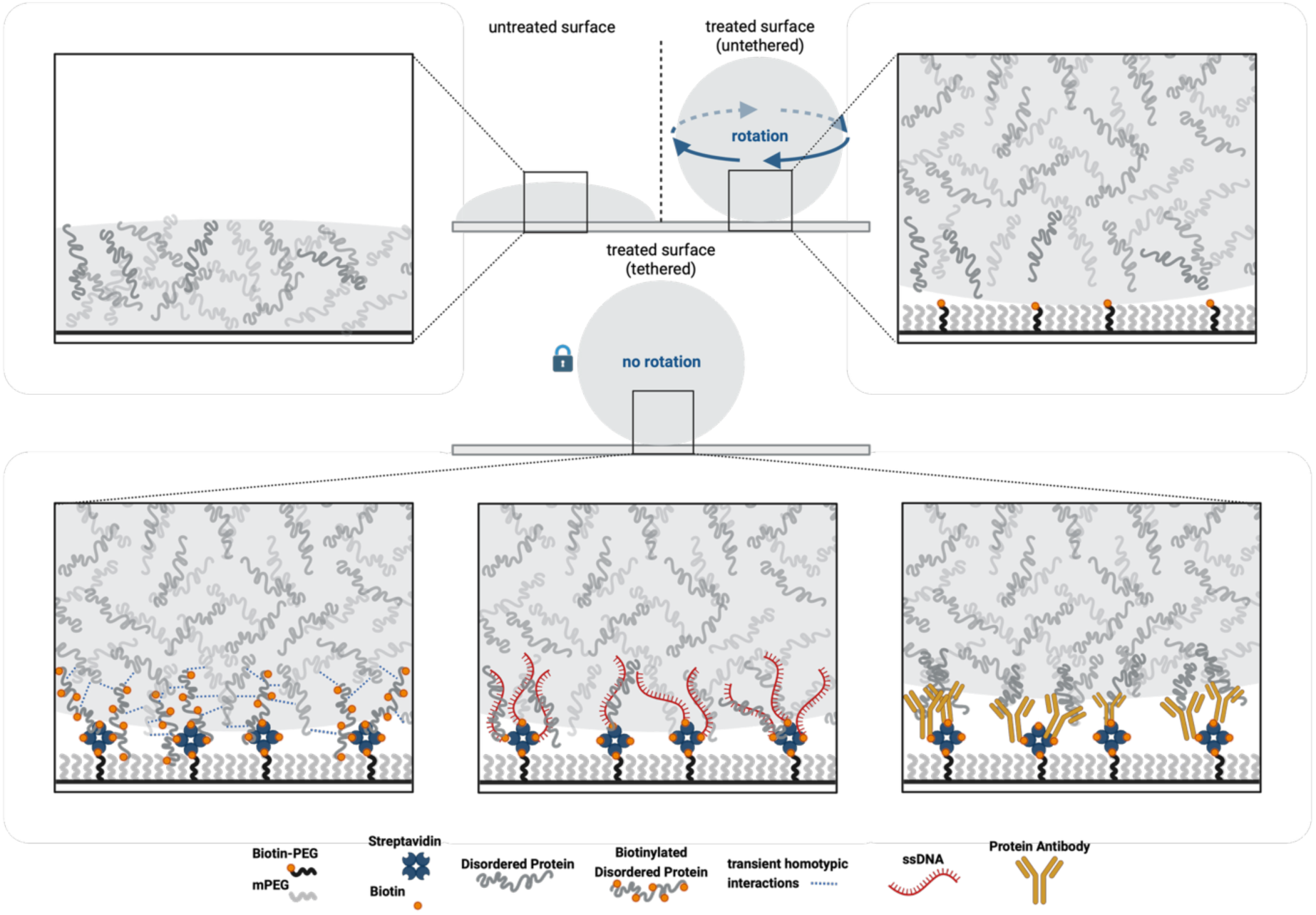

## Introduction

Biomolecular condensates, also known as membraneless organelles, have emerged as fundamental organizers of cellular biochemistry [1,2], orchestrating the spatial and temporal distribution of proteins, RNAs, and other macromolecules without the need for membrane boundaries [3–5]. These dynamic assemblies form through liquid-liquid phase separation, enabling the compartmentalization of biochemical reactions and the control of cellular processes, including RNA metabolism, signal transduction, stress response, and genome organization [4,6–8]. Notable examples include nucleoli, stress granules, P-bodies [9], Cajal bodies [10], and germ granules, each contributing to critical aspects of gene expression, ribonucleoprotein complex assembly [11], and cellular adaptation to environmental cues [2].

To probe the internal organization and molecular dynamics within condensates, super-resolved position determination and single-molecule diffusion measurements have become indispensable tools [12–17]. Techniques such as single-molecule tracking (SMT) and super-resolution fluorescence microscopy allow researchers to quantify the mobility, interactions, and spatial distribution of individual molecules within the dense and heterogeneous environment of condensates. These approaches provide critical insights into the physical properties and functional mechanisms governing condensate behavior and are often performed in vitro.

However, a significant and often underappreciated challenge in these in vitro studies arises from the intrinsic Brownian motion of condensates in vitro. As colloidal entities, biomolecular condensates exhibit translational and rotational movements driven by thermal fluctuations when placed on untreated or inadequately passivated surfaces [18–20]. One potentially contributing such phenomenon is thermal Marangoni convection, which can arise from laser-induced heating in phase separated binary mixtures [21,22], although it has not yet been studied extensively in biomolecular condensates. Such overlaid, intrinsic condensate motions have the potential to introduce substantial artifacts into diffusion measurements, possibly leading to systematic over- or underestimation of molecular mobility and confounding the interpretation of single-molecule trajectories. In vitro studies have yet to address that poorly immobilized condensates can compromise the accuracy of diffusion data analysis, particularly if nanometer spatial resolution and millisecond temporal resolution are desired, where whole-condensate movements can be mistaken for intra-condensate molecular diffusion.

Fused in Sarcoma (FUS) protein has become a widely used model for studying biomolecular condensates due to its robust phase-separation behavior and its direct pathological relevance [23–26]. FUS is an RNA-binding protein implicated in the formation of stress granules and other nuclear and cytoplasmic condensates [27]. Mutations and aberrant phase behavior of FUS are linked to neurodegenerative disorders such as amyotrophic lateral sclerosis (ALS) and frontotemporal dementia (FTD), where abnormal condensate dynamics and aggregation contribute to disease progression [24,26]. Thus, accurate measurement of molecular diffusion and organization within FUS condensates is critical for understanding both fundamental biophysical principles and disease mechanisms.

In this study, we address the challenge of condensate motion artifacts by developing and systematically evaluating three surface-tethering strategies—using biotinylated DNA, protein, or antibody tethers—to immobilize FUS condensates on passivated glass surfaces. Our findings reveal that controlled surface passivation and tethering are critical for maintaining the native spherical morphology [28] and structural stability of FUS protein condensates. We employ single-molecule tracking and diffusion analysis to quantify the impact of tethering on both condensate stability and the precision of intra-condensate diffusion measurements. Our results demonstrate that surface tethering effectively suppresses whole-condensate Brownian motion, eliminates measurement artifacts, and preserves the native liquid-like properties essential for biological function. We further show that the necessity of tethering depends on the timescale of molecular diffusion relative to condensate motion, with implications for experimental design and data interpretation. By providing a robust methodological framework, our work enables more accurate and reproducible investigations into the molecular dynamics of membraneless organelles, advancing our understanding of their roles in cellular organization, regulation, and disease. Collectively, our results establish surface tethering as a necessary and tunable approach for the accurate quantification of intra-condensate molecular dynamics, with broad implications for the study of membraneless organelles and their roles in health and disease.

## Results

### Surface passivation and tethering govern morphology and stability of FUS protein condensates

Recently, evidence has emerged supporting a critical impact of surface treatment and tethering strategies on the morphology and behavior of biomolecular condensates [28,29]. To further test this notion, we used confocal microscopy to image condensates across multiple z-slices and assess their three-dimensional shape under various surface treatment conditions. On untreated bare glass surfaces, condensates exhibited substantial wetting interactions, spreading flatly and losing their roundness with a mean circularity (measured in two dimensions) of 0.69 ± 0.02 and a notably low object count of n = 120 (Figure 1a,b). In contrast, methylated PEG(henceforth simply “PEG”)-treated surfaces effectively restored physiologically relevant spherical morphology, significantly improving circularity to 0.78 ± 0.01 (n = 450), confirming the efficacy of PEG in preventing deleterious surface wetting interactions.

**Figure 1.**
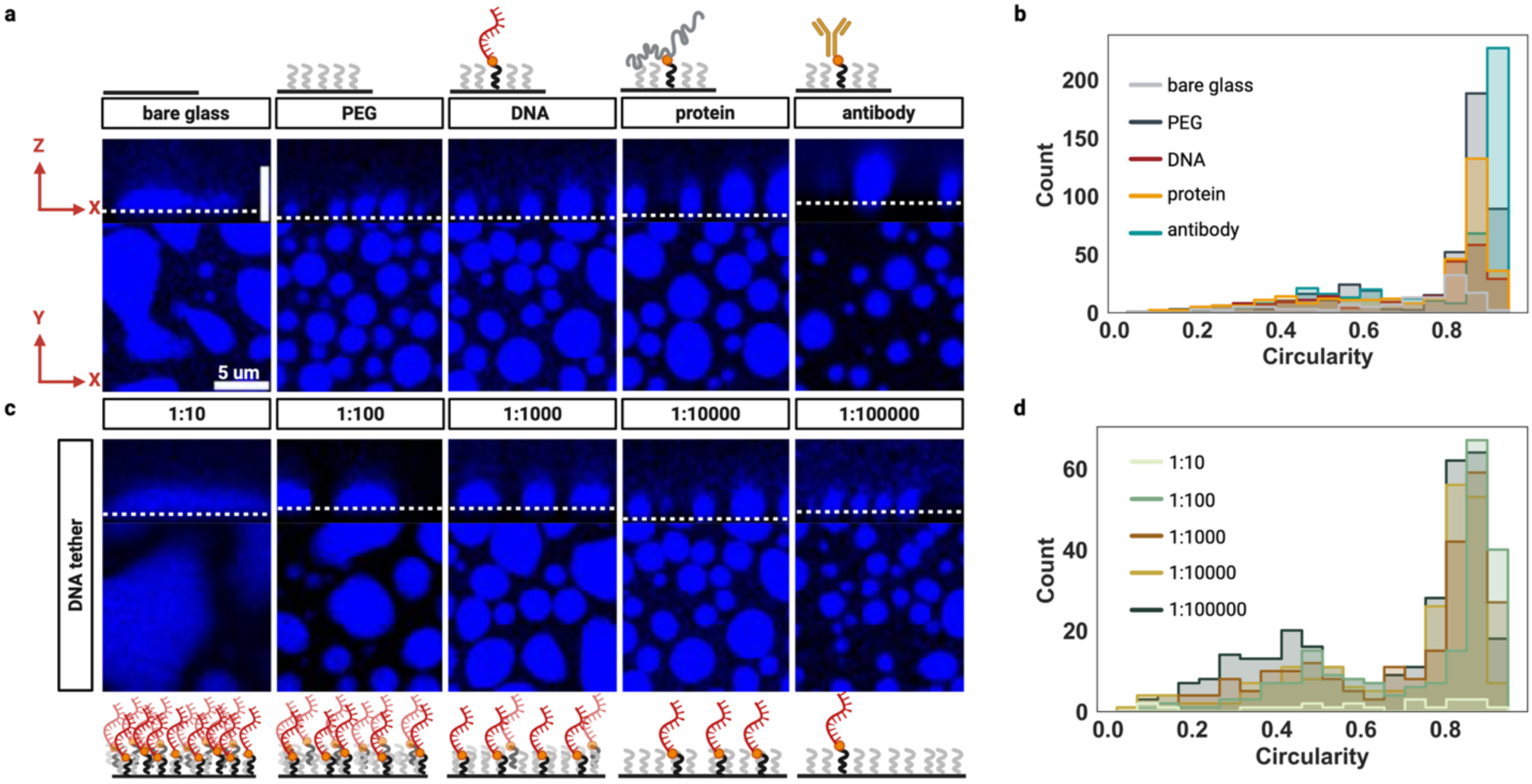
Surface modification and tethering strategies control biomolecular condensate morphology and stability. **a |** Confocal microscopy images showing the effects of different surface treatments on condensate morphology. Top row: schematic illustrations of surface functionalization-bare glass, PEG, DNA tether, protein tether, and antibody tether. Middle row: xz cross-sectional views reveal that condensates on bare glass exhibit extensive wetting and spreading, while PEGylation restores spherical morphology by preventing surface interactions. DNA, protein, and antibody tethers further stabilize condensates, maintaining roundness with slight deviations from perfect sphericity. Bottom row: corresponding xy images of condensate populations for each condition. **b |** Distributions of condensate circularity corresponding to identifiable objects in **a**. **c |** Tuning condensate morphology by varying the density of DNA tethers. Left to right: decreasing the ratio of biotinylated DNA PEG to PEG from 1:10 to 1:100000 progressively improves condensate roundness and reduces spreading, with the lowest tether density (1:100000) fully recovering spherical morphology and reducing condensate size. Schematics below illustrate the relative density of DNA tethers for each condition. Scale bar: 5 μm. **d |** Distributions of condensate circularity corresponding to identifiable objects in **c**.

Next, while our PEG passivation successfully recovered spherical condensate morphology, random Brownian whole-condensate motion emerged as a challenge due to the colloid-like properties of condensates [18,20] (Figure S1). To address this impediment to accurate intra-condensate measurements, three biotinylated tethering strategies were evaluated: DNA containing FUS binding sites, FUS protein, and anti-FUS antibody (Figure 1a).

All three tethering approaches successfully maintained condensate roundness while reducing Brownian motion. Notably, the anti-FUS antibody tethering achieved the highest mean circularity (0.80 ± 0.01, n = 437), representing a modest but measurable improvement over untethered PEG-treated surfaces. Biotin-FUS protein tethering yielded intermediate circularity values (0.74 ± 0.01, n = 334), while DNA tethering produced slightly lower circularity (0.72 ± 0.01, n = 231) compared to untethered PEG controls (Figure 1b). To systematically investigate the relationship between tethering density and condensate morphology, DNA tethering densities were tested by varying the proportion of biotin-PEG molecules relative to unmodified PEG molecules from 1:10 to 1:100,000 (Figure 1c,d).

At the highest tethering density (1:10 ratio), condensates exhibited severely compromised circularity (0.60 ± 0.05, n = 23)―resembling bare glass conditions―due to excessive surface wetting interactions caused by the dense lawn of attractive DNA molecules (Figure 1d). As tethering density decreased, condensate morphology progressively improved, with optimal circularity achieved at intermediate ratios. The 1:100 ratio produced the highest mean circularity (0.74 ± 0.01, n = 203), while further dilution to 1:1,000 (0.72 ± 0.01, n = 231), 1:10,000 (0.70 ± 0.02, n = 223), and 1:100,000 (0.66 ± 0.01, n = 316) showed gradual decreases in mean circularity, possibly due to weak surface binding (Figure 1d).

Interestingly, while mean circularity peaked at the 1:100 ratio, analysis of the circularity distribution revealed that the 1:100,000 ratio produced the highest frequency of highly circular condensates (circularity > 0.8), suggesting that optimal tethering conditions depend on the specific experimental requirements for condensate shape uniformity versus mean circularity (Figure 1d).

This tunable tethering system demonstrates that both tether type and density can be optimized to balance condensate sphericity and motility reduction for specific experimental applications. Our quantitative analysis reveals that surface modification strategies must be carefully calibrated to achieve reproducible imaging conditions while preserving the native liquid-like properties of biomolecular condensates as indicated by their sphericity.

### Tethering FUS condensates suppresses Brownian dynamics

To investigate the impact of tethering on Brownian dynamics of individual condensates, we developed an experimental approach to quantitatively assess thermal-induced motions using fluorescent microsphere tracers. Recent theoretical work has demonstrated that Brownian motion of droplets at submicron scales involves the complex interplay of intermolecular diffusion, surface tension, and thermal composition noise [19], which can be overlaid by thermal Marangoni convection [21,22]. Our protein condensate system models a liquid-liquid phase separation environment where such Brownian dynamics principles apply.

Based on the optimized DNA tether density achieved with a 1:1000 biotin-PEG to PEG ratio, we employed 200 nm fluorescent microspheres as internal tracers in FUS condensates due to their optimal balance of brightness for reliable frame-to-frame localization and sufficiently slow diffusion kinetics relative to condensate motion. Imaging parameters were carefully optimized: at 20 ms intervals, microsphere movement remained within static localization error limits, while 100 ms intervals allowed detectable displacement beyond the frame-to-frame detection threshold. Time-lapse imaging over 100 frames at 100 ms intervals provided sufficient temporal resolution to capture Brownian dynamics while maintaining spatial precision for individual particle tracking.

Qualitative analysis of the resulting microsphere trajectories revealed markedly different movement patterns between tethered and untethered conditions (Figure 2). In untethered condensates, microspheres exhibited highly coordinated displacement patterns, with trajectories color-coded by frame number demonstrating collective directional shifts characteristic of whole-condensate Brownian motion, most likely surface-induced thermal capillary wave motions (Figure 2a). This coordinated motion reflects the stochastic dynamics of the entire condensate as a colloidal particle undergoing thermal-driven Brownian motion in solution.

**Figure 2.**
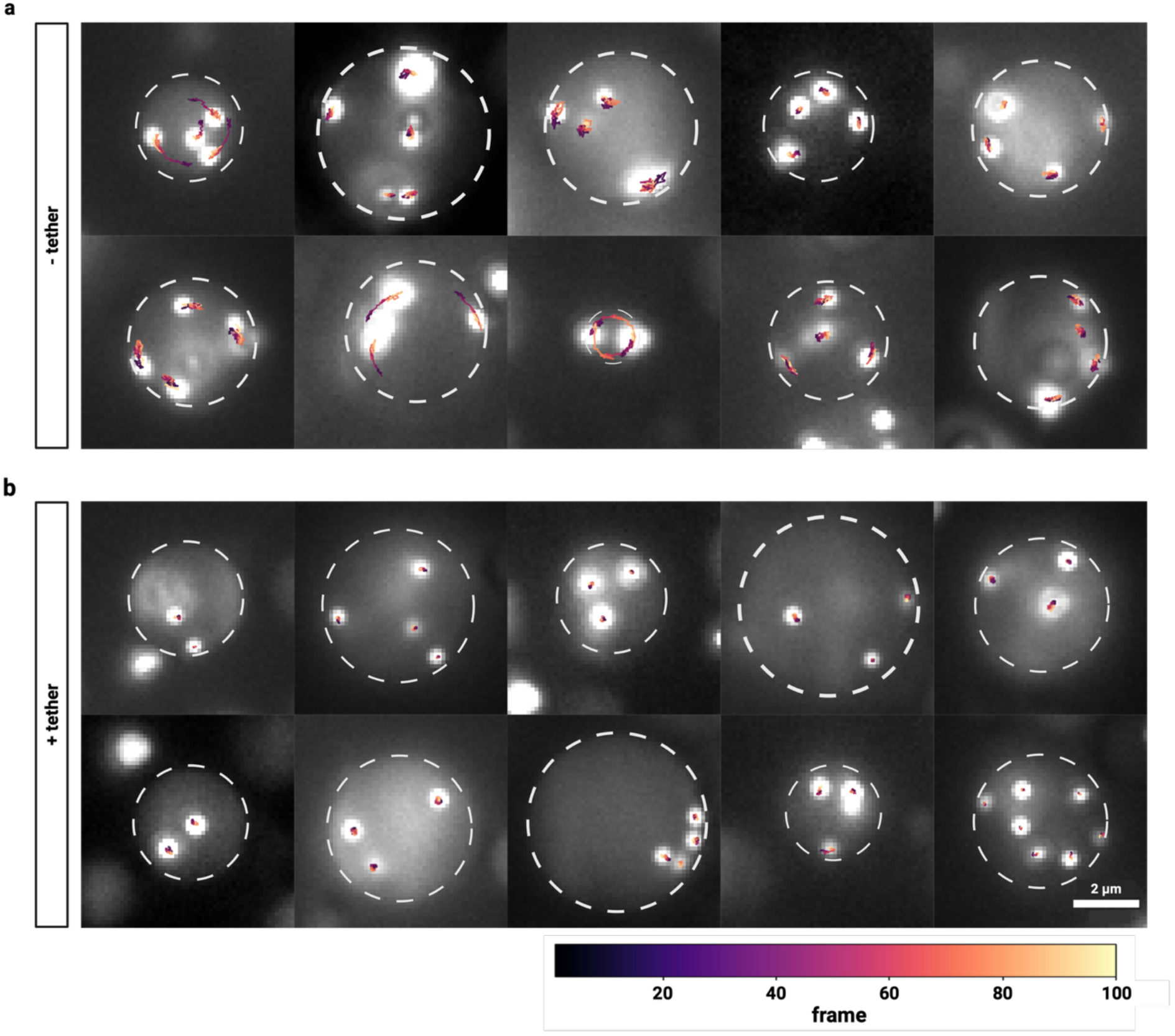
Tethering stabilizes biomolecular condensates against Brownian motion―most likely surface-induced thermal capillary wave motions. **a-b |** Single-molecule tracking of 200nm fluorescent microspheres within (a) untethered and (b) tethered biomolecular condensates. Ten representative condensates are shown for each condition. White dashed circles indicate condensate boundaries. Colored trajectories show microsphere positions across 100 frames (10 seconds), with colors indicating frame number according to the scale bar below. In untethered condensates (a), microspheres display coordinated movements with arc-like trajectories characteristic of condensate rotation driven by thermal capillary waves. In tethered condensates (b), microspheres exhibit random, uncorrelated diffusion patterns confined within stable condensates. Scale bar: 2 μm.

The observed collective microsphere movements indicate that the condensate undergoes both translational and rotational Brownian motion as predicted by established theory for spherical particles in viscous media (Figure 2a, S1). The coherent nature of these movements across multiple microspheres within individual condensates confirms that thermal fluctuations drive whole-condensate displacement rather than individual microsphere diffusion within a static condensate matrix.

In stark contrast, tethered condensates showed fundamentally different microsphere behavior (Figure 2b). Microsphere trajectories appeared incoherent and randomized, lacking the collective directional patterns observed in untethered conditions. This transition from coherent whole-condensate motion to localized microsphere diffusion indicates that going from untethered to surface-tethered effectively suppresses the thermal Brownian dynamics of the condensate as a colloidal entity while preserving local diffusive processes within the condensate interior.

### Tethering eliminates rotational coherence and reduces overall movement fluctuations within condensates

A quantitative analysis of angular displacement patterns revealed dramatic differences in rotational dynamics between tethered and untethered condensates (Figure 3a). Tracking the frame-to-frame fluctuations of microsphere localization revealed a dramatic increase in fluctuations for the untethered condition (Figure 3b). Further, by analyzing the angular displacement of microspheres relative to the condensate center, we calculated a maximum rotational amplitude of 72.95° ± 5.55° for the untethered condition (Figure 3b). These extensive rotational movements were largely eliminated in the tethered condition, where maximum angular displacement was reduced to 5.98° ± 3.04° (Figure 3b).

**Figure 3.**
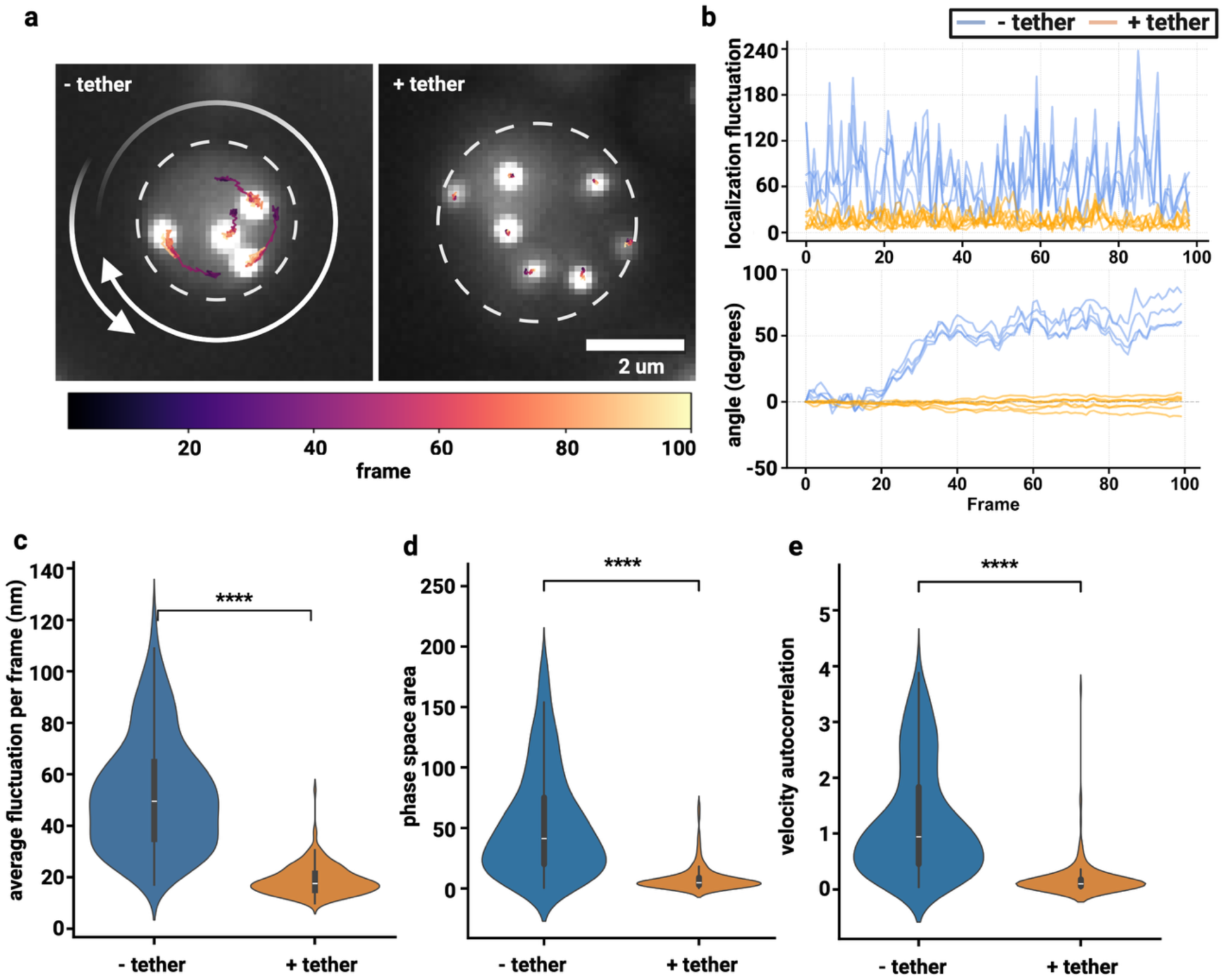
Untethered condensates display increased fluctuations and rotational coherence compared to tethered condensates. **a |** representative condensate for untethered and tethered conditions. Colored trajectories show microsphere positions across 100 frames (10 seconds), with colors indicating frame number according to the scale bar below. **b |** Frame-to-frame localization fluctuations and angle-of-rotation for the given condensates in **(a)** untethered (blue) and tethered (orange). **c |** Ensemble frame-to-frame localization fluctuations for all condensates in each condition. **d |** Scatter plot of phase space area as a function of angular velocity-acceleration correlations. N_untethered_ = 320, N_tethered_ = 183. Statistics annotation: Mann-Whitney U, ns: 0.05 < p <= 1 or Cliff’s |δ| < 0.147, *: 0.01 < p <= 0.05, **: 0.001 < p <= 0.01, ***: 0.0001 < p <= 0.001, ****: p <= 0.0001.

Ensemble localization fluctuation analysis demonstrated that untethered condensates exhibited maximum translational fluctuations of 153.7 ± 3.7 nm and average fluctuations of 52.4 ± 1.2 nm per frame (Figures 3c, S5a). In stark contrast, tethered condensates showed substantially reduced translational fluctuations with maximum values of 52.4 ± 1.2 nm and average fluctuations of only 18.9 ± 0.4 nm per frame, representing a three-fold reduction in maximum fluctuation amplitude and a 2.8-fold decrease in average fluctuation magnitude (Figures 3c, S5a).

Phase space area analysis provided quantitative evidence for the constrained dynamics observed in tethered condensates. Untethered condensates occupied significantly larger regions of velocity-acceleration phase space (53.7 ± 2.4 rad²/s³) compared to tethered condensates (8.2 ± 0.8 rad²/s³; p = 1.74e-51, Cliff’s δ = 0.81, large effect; Figure 3d). This 6.5-fold reduction in accessible phase space area demonstrates that tethering fundamentally restricts the range of translational motions available to the condensate as a whole. Velocity autocorrelation analysis, which quantifies the persistence of rotational motion, showed similar dramatic differences. Untethered condensates exhibited autocorrelation values of 1.23 ± 0.05 compared to 0.19 ± 0.03 in tethered condensates (p = 4.92e-55, Cliff’s δ = 0.84, large effect), indicating that rotational motions in untethered condensates are highly persistent and coordinated over time (Figure 3e). These trends hold consistent across the 100 ms and 20 ms imaging frequencies (Figure S2).

### Tethering eliminates condensate motion artifacts to enable precise diffusion measurements of mRNA guest molecules

To comprehensively assess how surface tethering affects the measurement of biologically relevant molecular dynamics, we tracked in vitro transcribed and fluorescently labeled FL mRNA within both tethered and untethered condensates. We imaged at 200 ms time resolution, the same time scale at which we observed condensate Brownian dynamics, to capture the slow diffusion of single AlexaFluor 647-labeled firefly luciferase (FL) mRNA guest molecules. This approach allowed direct comparison of diffusion measurements for a typical structured RNA, which exhibits more complex intra-condensate behavior than the inert microsphere tracer used in our earlier experiments.

Localization fluctuation analysis confirmed that tethering significantly reduces measurement variability for biomolecular tracking. Untethered condensates exhibited maximum fluctuations of 242.2 ± 5.7 nm and average fluctuations of 106.8 ± 2.7 nm per frame, compared to tethered condensates that showed reduced maximum fluctuations of 193.2 ± 5.1 nm and average fluctuations of 70.1 ± 2.03 nm (p = 4.1e-29; Figures 4a, S6a). This 1.5-fold reduction in localization variability demonstrates that tethering provides a more stable experimental platform for accurate single-molecule measurements by minimizing artifacts from whole-condensate Brownian motion.

**Figure 4.**
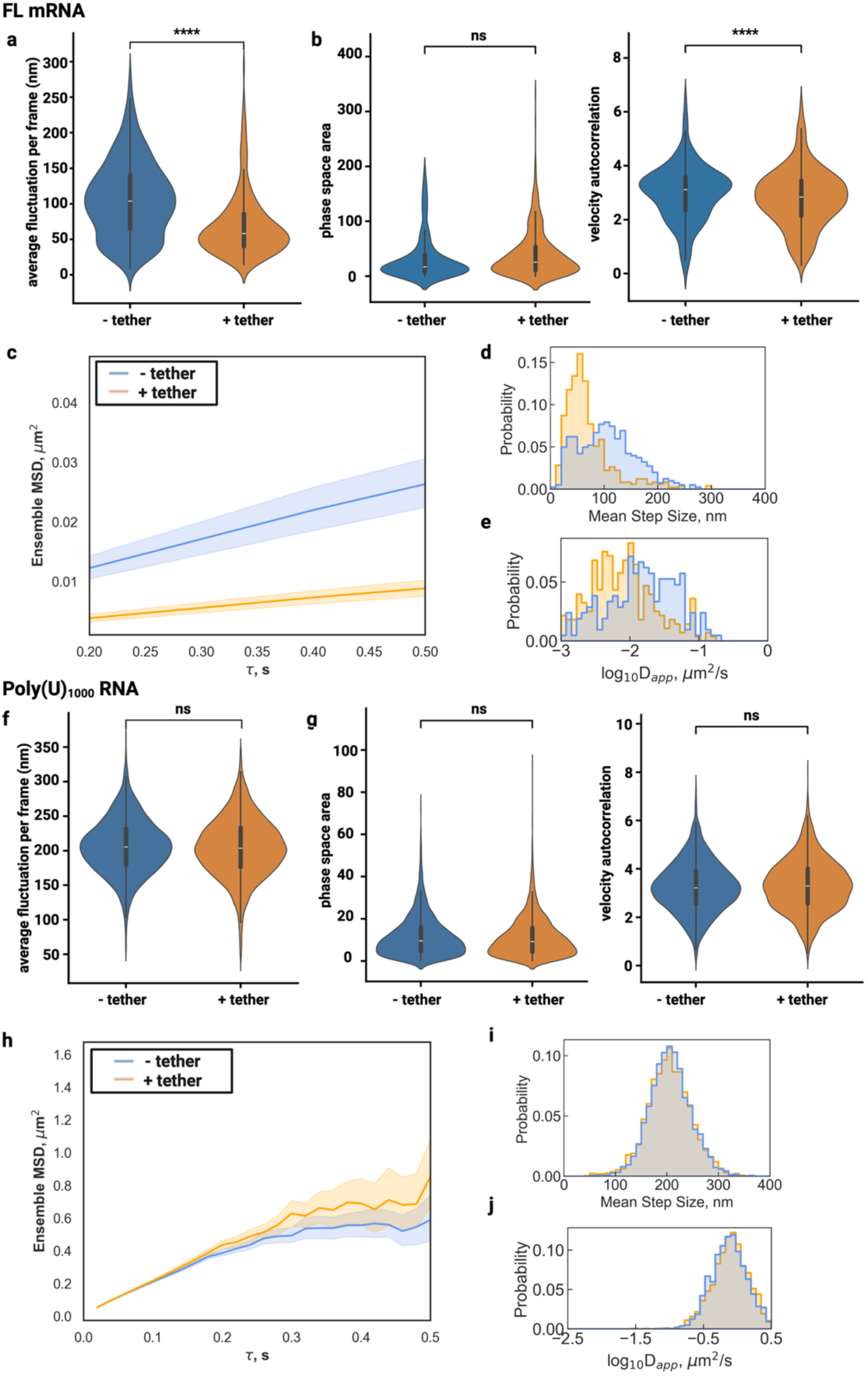
Tethering preserves intra-condensate diffusion dynamics while reducing condensate motion artifacts. **a |** Ensemble frame-to-frame localization fluctuations for all condensates in each condition; untethered (blue) and tethered (orange). N > 100. **b |** Violin of phase space area as a function of angular velocity-acceleration correlations and violin plot of velocity autocorrelation. N_untethered_ = 403, N_tethered_ = 493. Statistics annotation: Mann-Whitney U, ns: 0.05 < p <= 1 or Cliff’s |δ| < 0.147, *: 0.01 < p <= 0.05, **: 0.001 < p <= 0.01, ***: 0.0001 < p <= 0.001, ****: p <= 0.0001. **c |** Line plots of ensemble-averaged MSD (μm^2^) for RNA molecules in tethered and untethered condensates as a function of lag time (τ, seconds). Shaded regions represent the standard error of the mean. **d |** Histogram distributions of mean step sizes (nm) per trajectory. **e |** Histogram distributions of log-transformed apparent diffusion coefficients (log_10_D_app_, μm^2^/s). **f |** Ensemble frame-to-frame localization fluctuations for all condensates in each condition; untethered (blue) and tethered (orange). **g |** Violin plot of phase space area as a function of angular velocity-acceleration correlations and violin plot of velocity autocorrelation. N_untethered_ = 3,469, N_tethered_ = 2,645. Statistics annotation: Mann-Whitney U, ns: 0.05 < p <= 1 or Cliff’s |δ| < 0.147, *: 0.01 < p <= 0.05, **: 0.001 < p <= 0.01, ***: 0.0001 < p <= 0.001, ****: p <= 0.0001. **h |** Line plots of ensemble-averaged MSD (μm^2^) for RNA molecules in tethered and untethered condensates as a function of lag time (τ, seconds). Shaded regions represent the standard error of the mean. **i |** Histogram distributions of mean step sizes (nm) per trajectory. **j |** Histogram distributions of log-transformed apparent diffusion coefficients (log_10_D_app_, μm^2^/s).

Angular dynamics analysis of the FL mRNA location trajectories also revealed significant differences between tethered and untethered conditions across kinematic parameters. Phase space area analysis showed a modest difference between conditions, with untethered condensates occupying 23.98 ± 1.61 rad²/s³ compared to 13.77 ± 0.74 rad²/s³ in tethered condensates (p = 1.75e-03, Cliff’s δ = 0.12, negligible effect; Figure 4b). This smaller effect size for FL mRNA compared to the dramatic differences observed with microsphere tracers suggests that the complex diffusion behavior of mRNA molecules may partially buffer against whole-condensate motion effects. Velocity autocorrelation showed higher values in untethered conditions (2.21 ± 0.06) versus tethered conditions (1.21 ± 0.05; p = 1.07e-31, Cliff’s δ = 0.45, medium effect; Figure 4b).

FL mRNA in tethered condensates showed consistently lower mean squared displacement across all time lags (Figure 4c). Step size analysis revealed the impact of condensate motion on apparent molecular displacement measurements. FL mRNA molecules in untethered condensates displayed significantly larger mean step sizes (106.13 ± 3.20 nm, n = 270) compared to tethered condensates (66.36 ± 2.10 nm, n = 368; Figure 4d). This 1.6-fold difference in step size directly translates to systematic overestimation of molecular mobility when condensate motion is not controlled.

Analysis of the apparent diffusion constant (D_app_) and anomalous diffusion component (α) revealed that tethering preserves the fundamental diffusion characteristics of mRNA within condensates. The α values were similar between conditions, with tethered condensates showing 0.80 ± 0.01 (n = 352) compared to 0.75 ± 0.02 (n = 278) in untethered condensates (Figure S3). These values, close to α = 1, confirm that mRNA diffusion within condensates remains predominantly Brownian regardless of tethering status, indicating that surface immobilization preserves the native liquid-like properties essential for condensate function. Apparent diffusion coefficient analysis quantified the magnitude of this measurement artifact. In untethered condensates, the mean log₁₀(D_app_) was -2.15 ± 0.04 log₁₀(μm²/s) (n = 378), compared to -2.42 ± 0.03 log₁₀(μm²/s) in tethered condensates (n = 476; Figure 4e). This difference represents approximately a 1.9-fold overestimation of diffusion coefficients when condensate Brownian motion is not eliminated, demonstrating how failure to account for whole-condensate movement can significantly confound single-molecule diffusion measurements. Similar concerns would also apply to bulk measurements such as fluorescence recovery after photobleaching (FRAP).

### Tethering is not necessary for measuring fast diffusion of unstructured RNA molecules

We next asked whether tethering condensates has a similar impact on the quantitative analysis of the diffusive behavior of unstructured, fluorescently labeled Poly(U)₁₀₀₀ RNA molecules compared to structured RNA. To capture the diffusion of Poly(U)₁₀₀₀, imaging had to be performed at an order of magnitude faster frequency of 20 ms compared with FL mRNA analyzed at 200 ms. Our findings contrast sharply with the dramatic differences in Brownian condensate dynamics observed for structured FL mRNA, providing insights into the temporal scales at which whole-condensate motion affects molecular tracking.

Frame-to-frame localization fluctuation analysis revealed remarkably strikingly similar behavior between tethered and untethered condensates containing guest Poly(U)₁₀₀₀ RNA molecules. Average fluctuations were virtually identical between conditions, with tethered condensates showing 204.5 ± 0.8 nm and untethered condensates exhibiting 206.3 ± 0.7 nm (Figure 4f). These results suggest that individual Poly(U)₁₀₀₀ molecules retain equivalent short-timescale mobility regardless of any overlaid condensate Brownian dynamics.

Comprehensive angular dynamics analysis confirmed the absence of significant differences between tethered and untethered conditions for Poly(U)₁₀₀₀ RNA tracking. Phase space area analysis revealed minimal differences (10.92 ± 0.17 rad²/s³ tethered versus 11.86 ± 0.17 rad²/s³ untethered, p = 2.33e-03, Cliff’s δ = 0.05, negligible effect; Figure 4g), with both distributions highly clustered and exploring limited phase space compared to the microsphere tracers and structured mRNA molecules analyzed in our previous experiments.

Analysis of molecular diffusion parameters demonstrated that tethering provides no measurement advantage for unstructured RNA molecules. MSD-*τ* plot residuals overlapped even at larger time lags, with untethered condensates displaying lower average values at larger time lags (Figure 4h). Mean step sizes were essentially identical between conditions (204.76 ± 0.93 nm for tethered, n = 1,947; 205.66 ± 0.76 nm for untethered, n = 2,587), with overlapping distributions (Figure 4i). The anomalous diffusion component (α) remained close to Brownian behavior in both conditions (0.86 ± 0.01 tethered versus 0.84 ± 0.01 untethered), confirming that the fundamental diffusion characteristics of unstructured RNA are preserved regardless of condensate stabilization status (Figure S3). Log-transformed apparent diffusion coefficients showed similarly negligible differences (-0.32 ± 0.01 log₁₀(μm²/s) tethered versus -0.33 ± 0.01 log₁₀(μm²/s) untethered; Figure 4j).

The minimal impact of tethering on Poly(U)₁₀₀₀ RNA measurements can be explained by the temporal scale separation between molecular diffusion and condensate Brownian dynamics. The frame-to-frame diffusion timescale of unstructured Poly(U)₁₀₀₀ RNA molecules (10-20 ms) is approximately one order of magnitude faster than the characteristic timescales of whole-condensate Brownian motion observed in previous experiments (100-200 ms). This temporal separation means that rapid RNA diffusion effectively dominates over slower condensate motions, rendering tethering unnecessary for accurate measurement of intrinsic molecular dynamics.

### Different tethering strategies offer tunable control over condensate stability and dynamics

To determine the optimal tethering approach for different experimental applications, we systematically compared our three distinct tethering methods—DNA containing FUS binding sites (n = 2,100), biotin-FUS protein (n = 3,221), and anti-FUS antibody (n = 2,779),—against untethered controls (n = 415) using FL mRNA as a representative structured molecule with complex diffusion behavior. This comprehensive comparison was imaged at 20 ms to capture all diffusion states. We evaluated the relative effectiveness of each approach in suppressing Brownian condensate dynamics while preserving native molecular diffusion characteristics.

Frame-to-frame localization fluctuation analysis revealed that all three tethering methods provide comparable stabilization effectiveness. Comparing each method revealed that anti-FUS antibody tethering (71.7 ± 0.7 nm avg), biotin-FUS protein tethering (74.9 ± 0.7 nm avg), and DNA tethering (75.0 ± 0.8 nm avg) display fluctuations with similar distributions (Figure 5a). The statistical comparisons between tethering methods revealed negligible effect sizes (Cliff’s δ = -0.07 to -0.08), confirming that each approach provides equivalent localization stability. Notably, these fluctuations are a marked increase from the untethered condition (52.3 ± 1.5 nm avg; Figure S4) affirming our earlier observation that the slowly diffusing structured mRNA is fluctuating on the same time scale as whole-condensate motions.

**Figure 5.**
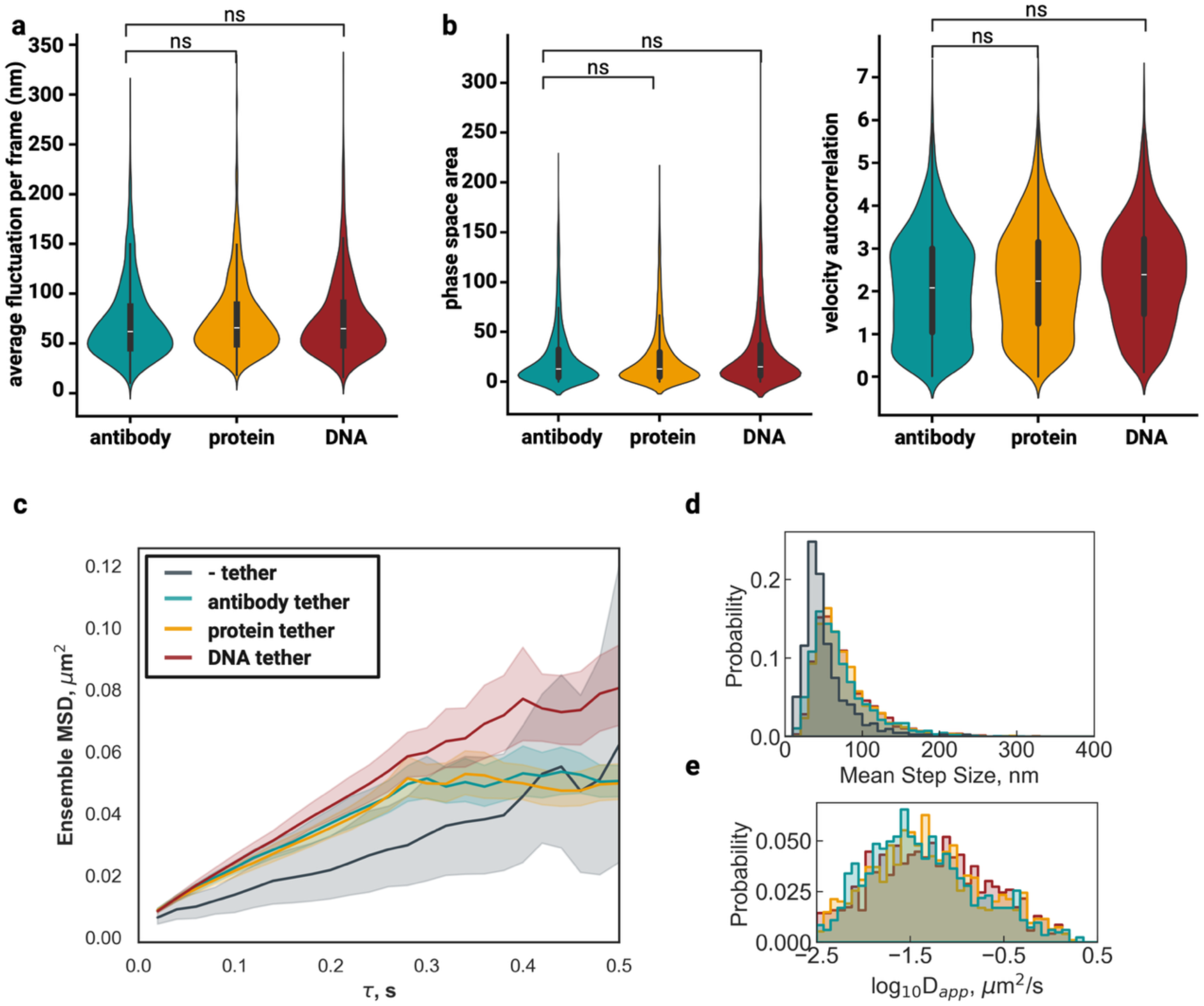
**a |** Ensemble frame-to-frame localization fluctuations for all condensates in each condition; antibody tethered (blue), protein tethered (yellow), and DNA tethered (red). Statistics annotation: Mann-Whitney U, ns: 0.05 < p <= 1 or Cliff’s |δ| < 0.147, *: 0.01 < p <= 0.05, **: 0.001 < p <= 0.01, ***: 0.0001 < p <= 0.001, ****: p <= 0.0001. **b |** Violin plot of phase space area as a function of angular velocity-acceleration correlations and violin plot of velocity autocorrelation. N_antibody tethered_ = 2,779, N_protein tethered_ = 3,221, N_DNA tethered_ = 2,100. **c |** Line plots of ensemble-averaged MSD (μm^2^) for RNA molecules in tethered condensates as a function of lag time (τ, seconds). Shaded regions represent the standard error of the mean. **d |** Histogram distributions of mean step sizes (nm) per trajectory. **e |** Histogram distributions of log-transformed apparent diffusion coefficients (log_10_D_app_, μm^2^/s).

Indeed, angular dynamics analysis demonstrated that all three tethering strategies effectively eliminate the large-scale Brownian motions observed in untethered condensates. Phase space area analysis showed dramatic reductions from untethered values (54.0 ± 3.8 rad²/s³) to all three tethered conditions: anti-FUS antibody (25.2 ± 0.6 rad²/s³), biotin-FUS protein (24.3 ± 0.5 rad²/s³), and DNA (28.5 ± 0.8 rad²/s³; Figure 5b). While statistically significant differences existed between tethering methods (p < 0.05), effect sizes remained negligible (Cliff’s δ ≤ 0.14), confirming that all approaches successfully suppress condensate Brownian dynamics. Similarly, negligible effect sizes were found for the velocity autocorrelation analysis between tethering methods (Figure 5b).

Despite equivalent condensate stabilization, tethering methods exhibited subtle, but measurable differences in molecular diffusion parameters. Notably, the MSD-*τ* relationship for the DNA tether condition showed consistently larger values across all time lags while antibody and protein tether overlapped entirely, and the untethered condition showed consistently lower values across all time lags (Figure 5c). Step size analysis also revealed systematic differences between approaches, with untethered condensates showing artificially reduced step sizes (46.8 ± 1.9 nm) due to whole-condensate motion artifacts masking true molecular displacement (Figure 5d). Among tethered conditions, anti-FUS antibody tethering produced the smallest step sizes (66.6 ± 0.9 nm), while protein (70.6 ± 0.9 nm) and DNA (71.6 ± 1.1 nm) tethering showed progressively larger values (Figure 5d).

The anomalous diffusion component (α) showed minimal variation between tethering methods (antibody: 0.61 ± 0.01, protein: 0.61 ± 0.01, DNA: 0.62 ± 0.01), all substantially higher than untethered controls (0.53 ± 0.02), confirming that tethering preserves normal diffusion characteristics while eliminating motion artifacts (Figure S4). Accordingly, apparent diffusion coefficient analysis revealed a consistent trend across tethering methods. Untethered condensates yielded artificially low apparent diffusion coefficients (-2.45 ± 0.04 log₁₀(μm²/s)) due to measurement artifacts from Brownian motion. DNA tethering produced the highest apparent diffusion coefficients (-1.77 ± 0.02 log₁₀(μm²/s)), followed by protein tethering (-1.81 ± 0.01 log₁₀(μm²/s)) and antibody tethering (-1.86 ± 0.02 log₁₀(μm²/s); Figure 5e).

The observed subtle differences in molecular diffusion parameters between tethering methods suggest distinct biophysical interactions at the condensate-surface interface. DNA tethering, which showed the highest apparent diffusion coefficients and largest step sizes, may create a more permissive internal environment for RNA diffusion through specific protein-nucleic acid interactions that slightly reduce local condensate viscoelasticity, that is, make it behave more fluid-like. Conversely, antibody tethering, which produced more constrained diffusion parameters, may introduce additional steric hindrance or alter condensate internal organization through protein-protein interactions.

### Necessity of tethering depends on condensate size and diffusion properties of the tracked molecules

To systematically assess the necessity of tethering for intra-condensate SMT beyond the conditions experimentally tested so far, we used Monte Carlo simulations to profile the deviation of SMT readouts in untethered versus tethered condensates across a broader parameter space (Figure 6). We generated SMT trajectories molecules with ground-truth diffusion metrics, including different diffusion coefficient (*D)* and anomalous coefficient (*α)* (Figure 6a). We added Brownian rotation of the condensate to molecular trajectories at each step of the simulation, based on rotational speed calculated from size (*R)* of the condensate, where molecules are at room temperature (*T)* with a viscosity (*η)* representing a typical protein solution. Finally, we extracted apparent diffusion coefficient (*D_app_*) and anomalous coefficient (*α_app_*) from simulated trajectories using the same MSD-ι− fitting method as above. To determine the effect of tethering, we performed pairs of tethered and untethered (without and with Brownian rotation added) simulations for each condition side-by-side to assess the impact of the lack of tethering on SMT metrics.

**Figure 6.**
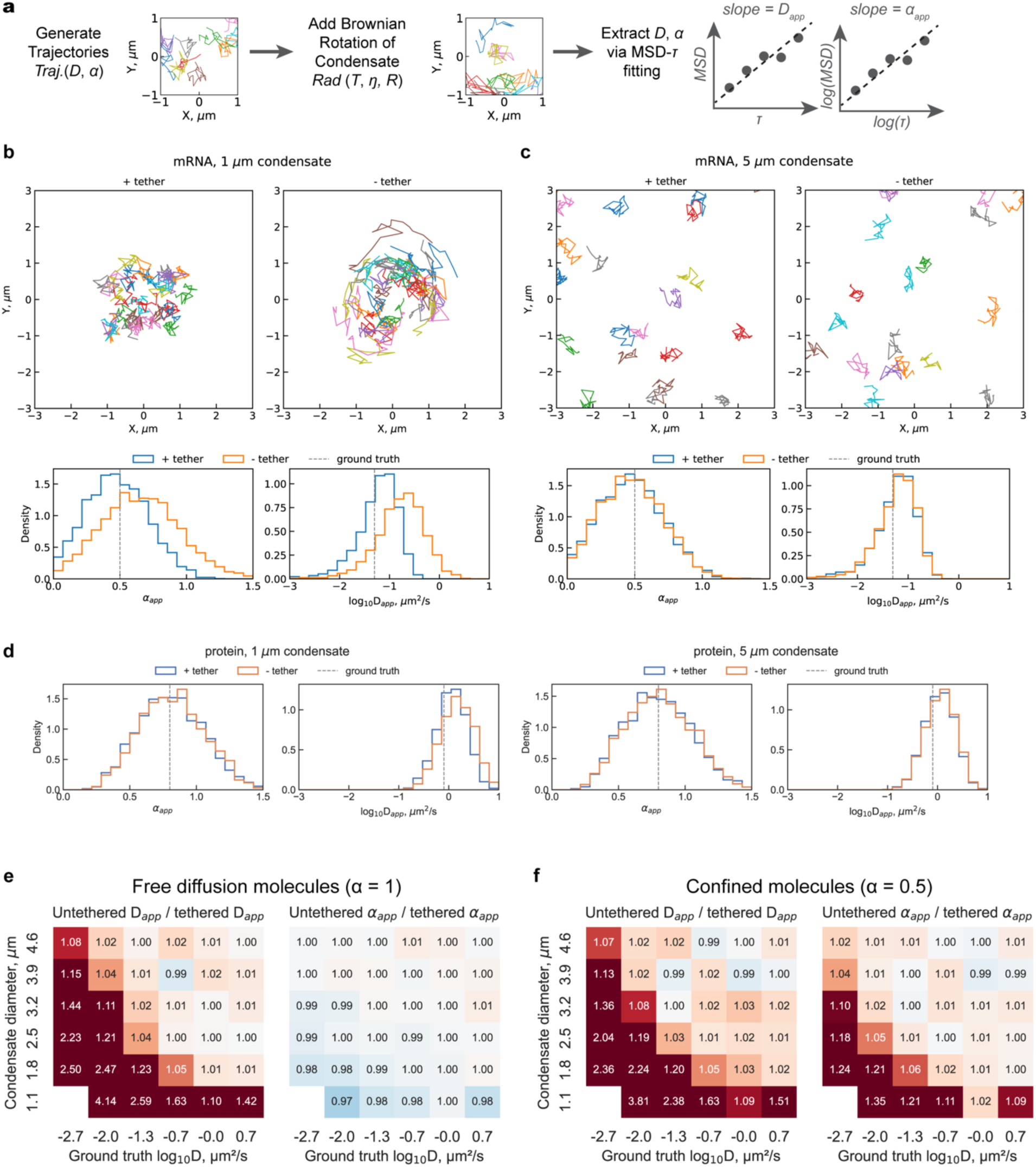
Simulations reveal parameter-dependent bias in intra-condensate SMT measurements due to lack of tethering. **a |** Schematic of the simulation workflow and parameters used for generating and analyzing SMT trajectories. **b-c |** Representative trajectories and distributions of *D_app_* and *α_app_* for a typically confined, slow-diffusing mRNA in a small (**b**) or large (**c**) condensate. **d |** Distributions of *D_app_* and *α_app_* for a typically free, fast-diffusing mRNA in an either small or large condensate, matching the size in **b-c**. **e-f |** Heatmaps showing the systematic bias introduced by the lack of tethering on *D_app_* and *α_app_* across physiological ranges of condensate size (0.3 – 5 μm) and ground-truth D (0.001 – 1 μm^2^/s) for free-diffusing (**e**) or confined (**f**) molecules. Color scale represents the ratio of apparent metrics measured in untethered versus tethered simulations: red indicates overestimation, blue indicates underestimation.

We first examined the impact of the lack of tethering on the SMT of confined, slow-diffusing, and free, fast-diffusing molecules inside condensates, representing mRNA and protein diffusion in condensates, respectively (Figure 6b-d) [29]. For confined, slow-diffusing molecules such as mRNAs within small condensates, the lack of tethering caused noticeable additional displacements in trajectories, resulting in overestimation of both *D_app_* and *α_app_* (Figure 6b). Notably, the distributions of *D_app_* and *α_app_* under the tethered condition were centered around the ground-truth *D* and *α*, confirming that our simulation system faithfully recapitulates the expected diffusion behavior of molecules (Figure 6b). The slight deviation from ground-truth *D* in the *D_app_* distribution is a known systematic error introduced by MSD-τ fitting [30], rather than the simulation itself. In contrast, the same confined, slow-diffusing molecules in larger condensates will not be affected, whether tethered or not (Figure 6c). Therefore, SMT metrics extracted from confined, slow-diffusing molecules are overestimated in small condensates but not in large condensates. For free, fast-diffusing molecules such as proteins, the distributions of *D_app_* and *α_app_* remain unchanged upon the lack of tethering, whether in small or large condensates (Figure 6d). This suggests that SMT measurements of free, fast-diffusing molecules are not biased by the lack of tethering. Taken together, simulation data on four biologically relevant diffusion scenarios showed that only confined, slow-diffusing molecules in small condensates are biased by the lack of tethering in the tracking experiments.

These observations from simulated data match the observations from the above experimental results, where condensate sizes are also around 1 μm, that tethering is necessary for slow-diffusing molecules (Figure 4a-e) but not fast-diffusing ones (Figure 4f-j) in physiological, small condensates.

To test if this principle is generally true, we next performed a broader parameter sweep using our simulation in physiological ranges of condensate size (0.3 – 5 μm) and ground-truth D (0.001 – 1 μm^2^/s) and compared the distribution of *D_app_* and *α_app_* under tethered versus untethered conditions (Figure 6e-f). The ratio between the same apparent SMT metric extracted from untethered over tethered conditions is plotted as a heatmap, where darker color indicates a stronger bias introduced by the lack of tethering (Figure 6e-f). For free-diffusing molecules (*α*=1), the slower the molecules are and the smaller the condensate they are in, the more overestimated their *D_app_* becomes (Figure 6e), suggesting that our observations under a couple combinations of parameters hold true in a broader parameter space. Interestingly, the estimation of *α_app_* in free-diffusing molecules is never affected by the lack of tethering, whether they diffuse slowly or rapidly (Figure 6e). This means the assignment of anomalous diffusion type will never be wrong for real free-diffusing molecules, regardless of tethering condition or diffusion rate. For confined molecules (*α*=0.5), the same conclusions remain for the *D_app_* but not *α_app_*, where *α_app_* would be overestimated for slow-diffusing confined molecules in small condensates (Figure 6f). An overestimated *α_app_* for confined molecules (*α*<1) can lead to a misassignment as free-diffusing molecules (*α*=1). Therefore, true confined molecules may be missed if appropriate tethering is not in place when the molecules diffuse slowly and are in a small condensate.

In summary, our simulation sweeping a broad parameter space validates our prior conclusion that the measurement of slow but not fast intra-condensate diffusion requires proper tethering. Moreover, it reveals two new principles for determining whether tethering is necessary for intra-condensate SMT: (1) For condensates larger than 2 μm in diameter, tethering generally does not affect the estimation of *D_app_* or *α_app_* (Figure 6e–f), because the Brownian rotation of condensates becomes negligible relative to the physiological *D* and *α* range for biomolecules; (2) Tethering is essential for the correct identification of confined diffusion in small condensates. Confined, slow-diffusing molecules such as mRNA [cite nanodomain paper] will be misassigned as free-diffusing molecules when the condensates are not properly tethered.

## Discussion

The results presented herein establish that surface tethering is a powerful and necessary strategy for eliminating artifacts arising from condensate Brownian dynamics in single-molecule diffusion measurements. By immobilizing condensates on passivated surfaces, we provide a stable reference frame that enables accurate quantification of intra-condensate molecular mobility, especially for large, structured mRNAs that diffuse slowly, revealing the true dynamic behavior of guest RNAs (and proteins) within phase-separated assemblies.

Our systematic comparison of three tethering strategies—DNA, protein, and antibody—demonstrates that all three approaches successfully suppress both translational and rotational condensate motion, as evidenced by dramatic reductions in localization fluctuations, angular displacement, and accessible phase space area (Figures 3, 4, 5). These findings directly address a longstanding phenomenon in the literature: untethered condensates, due to their colloidal nature, undergo stochastic movements [18,19,31–35] that can be misinterpreted in SMT measurements as reduced or enhanced molecular diffusion (Figure 4). Our results provide direct experimental evidence that such artifacts are prevalent in untethered systems and must be controlled for rigorous biophysical analysis.

Our comparative analysis of tethering methods further reveals that the choice of tether can subtly influence internal molecular mobility. DNA tethers facilitate enhanced RNA mobility, potentially due to specific protein-nucleic acid interactions that alter local viscoelastic properties [11,36,37], while antibody tethers slightly restrict molecular movement, possibly through steric effects or altered condensate organization (Figure 5). These findings underscore the importance of optimizing tethering conditions to balance condensate stability, preservation of native dynamics, and compatibility with the biological question at hand.

Importantly, our experiments and simulations demonstrate that the impact of tethering is context dependent (Figure 4, 6) and leads to the following practical guideline: (1) Appropriate tethering is essential only for slow-diffusing, confined molecules in small condensates, as neglecting tethering under these conditions leads to significant overestimation of diffusion parameters and potential misclassification of diffusion modes; (2) For fast-diffusing or unconfined molecules and for condensates larger than 2 μm, tethering is generally unnecessary. Thus, the necessity of condensate stabilization should be evaluated based on the dynamical properties of the molecules and the size of the condensates under investigation, providing a robust framework for the design and interpretation of intra-condensate SMT experiments.

The implications of this work extend beyond methodological rigor. In vivo, many membraneless organelles are anchored to cytoskeletal elements [38,39], chromatin [6,40], or membrane surfaces [17,41], restricting their motion and enabling precise spatial regulation of biochemical processes. Our tethering approach provides a more physiologically relevant in vitro platform that mimics this confinement, facilitating the study of molecular exchange, internal reorganization, and response to perturbations under controlled conditions.

Finally, we discovered a marked, ∼2 orders of magnitude increase―from D_app_ = -2.42 ± 0.03 μm²/s to D_app_ = -0.32 ± 0.01 log₁₀(μm²/s)―in mean apparent diffusion constant from the structured, ∼1,500 nt firefly luciferase mRNA to the unstructured Poly(U)₁₀₀₀, ∼1,000 nt in length. This difference stands in stark contrast to the ∼24% increase in D_app_ expected for the smaller Poly(U)₁₀₀₀ RNA based on its ∼1.5× lower molecular mass, assuming random coil structures. This finding suggests that the intrinsic secondary structure of an mRNA may play a role in slowing down diffusion, possibly due to its capacity to establish the extensive intermolecular base pairing proposed to play a role in the formation of RNA-protein condensates [42,43].

In summary, our study provides a robust framework for the quantitative analysis of molecular diffusion dynamics within biomolecular condensates. By eliminating motion artifacts through tunable surface tethering, we enable more accurate and reproducible investigations into the physical principles and biological functions of membraneless organelles.

## Materials & Methods

### Protein and RNA Purification and Labeling

MBP-FUS fusion protein was expressed in *E. coli* BL21(DE3) cells transformed with the MBP-FUS_FL_WT was a gift from Nicolas Fawzi (Addgene plasmid # 98651) [44]. A single colony was inoculated into 2 mL LB medium containing 100 µg/mL ampicillin and grown overnight at 37°C with shaking (160 rpm). This pre-culture was diluted 1:20 into 50 mL fresh LB-ampicillin and incubated under identical conditions to ensure robust cell viability. The entire pre-culture was then transferred into 1 L of LB-ampicillin (final volume 800 mL) and grown at 37°C until reaching mid-log phase (OD_600_ = 0.6–0.8). To enhance soluble protein production, cultures were cooled to 16°C (pre-chilled shaker) before induction with 0.5 mM IPTG. Expression proceeded overnight at 16°C with shaking (160 rpm), a temperature regime optimized to balance protein yield and solubility. Cells were harvested by centrifugation (6,500 × g, 7 min, 4°C), washed with 0.5× PBS, and stored at -80°C. Expression efficiency was confirmed via SDS-PAGE by comparing pre- and post-induction samples, with successful MBP-FUS production indicated by a dominant band at the expected molecular weight (∼98 kDa; Figure S5). The MBP-FUS fusion protein was purified using sequential affinity chromatography. First, amylose resin affinity chromatography captured the maltose-binding protein (MBP) tag, with optimal fractions selected for further processing. The eluate was then applied to a HisTrap HP column pre-equilibrated with MBP-HisWash buffer (500 mM imidazole), and fractions containing the His-tagged fusion protein were collected. To remove imidazole, the pooled fractions were concentrated via three parallel ultrafiltration columns (10 kDa MWCO), each diluted with 15 mL of imidazole-free MBP-Normal Wash buffer and re-concentrated. This diafiltration process was repeated twice, reducing imidazole concentration 100-fold to <5 mM. The final MBP-FUS product (6 mL) had an absorbance (A_280_) of 1.3, corresponding to a concentration of 0.92 mg/mL (9.4 µM) using an extinction coefficient of 139,720 M^-1^cm^-1^. For tag removal, the fusion protein was treated with TEV protease (Millipore Sigma), yielding cleaved MBP and untagged FUS (Figure S5).

Full-length human FUS was expressed in E. coli BL21(DE3) using pET-hFUS [23,45] (a gift from the Ishihama group) and culture was grown to OD_600_=1.0 at 37°C and induced with 1 mM isopropyl β-D-1-thiogalactopyranoside (IPTG). After 6 h of growth at 37°C, cells were harvested by centrifugation and purified following an established protocol [23,29,45] via sequential ion-exchange chromatography (Cytiva HiTrap Q, Capto S, SP HP columns) under denaturing conditions (6 M urea). Refolding was performed in a β-cyclodextran (BCD) containing buffer to prevent aggregation, with a two-step dialysis process (CAPSO/arginine buffer followed by HEPES/NaCl/BCD buffer). Purity was confirmed by SDS-PAGE (Figure S5). AlexaFluor 488 NHS Ester (Click Chemistry Tools) was used to label FUS (20:1 dye:protein ratio) post-refolding, followed by buffer exchange to remove unbound dye. Labeling efficiency was quantified via absorbance (NanoDrop).

AlexaFluor 647-labeled firefly luciferase (FL) mRNA was synthesized using an established protocol [46,47] by in vitro transcription (T7 promoter), followed by enzymatic capping (NEB), sequential polyadenylation with 2’-azido-dATP (Jena Biosciences) and rATP [46,47], and click-chemistry conjugation to sDIBO-alkyne dye (Thermo Fisher) (Figure S6). Similarly, polyuridylic acid (poly(U)) potassium salt (Sigma-Aldrich) underwent gel purification of the ∼1000 nt fraction, followed by an initial incorporation of 2’-azido-2’-dATP (Jena Biosciences) followed by polyuridylation with poly(U) polymerase (NEB), employing rUTP [46,47]. Finally, the same click-chemistry reaction with Alexa Flour 647 sDIBO alkyne (Thermo Fisher) was conducted to produce fluorophore-conjugated poly(U) RNA. Free dye was removed from both RNAs by multiple rounds of washing of the ethanol-precipitated purified RNA pellet, which was confirmed by denaturing PAGE (Figure S6).

### Surface Treatment

Glass coverslips were functionalized with (3-Aminopropyl)triethoxysilane (Sigma-Aldrich), mPEG-SVA (NEB), and biotin-PEG-SVA (NEB; 10,000:1 ratio unless specified otherwise), followed by DST treatment [48,49]. Surface treatment was completed by using a PEG-biotin:streptavidin:tether-biotin system (tether: DNA FUS binding site, FUS protein, or anti-FUS antibody) we developed. Biotinylated DNA FUS binding site (5’-/5Bioag/TCCCCGT(6x)-3’, IDT) was selected as a tether due to established interactions with FUS [50]. The biotinylation of protein and antibody tethers was achieved by reacting purified FUS protein or antibody (BioLegend) with Biotin-NHS ester (Sigma-Aldrich) in dimethylformamide, achieving a 1:5 molar ratio of protein-to-biotin. The reaction mixture was maintained at pH 8.5 to optimize NHS-ester reactivity and incubated for 1 hour in the dark to prevent photodegradation. Unconjugated biotin was removed through two cycles of centrifugal filtration using 10 kDa Amicon Ultra filters (Sigma-Aldrich) at 4,000 × g (swinging bucket rotor), with buffer exchange to physiological pH 7.4 during resuspension. The purified biotinylated FUS was aliquoted into 10 µL volumes and stored at -80°C.

### Condensate Assembly

A 10x phase separation buffer containing 200 mM tris(hydroxymethyl)aminomethane (Tris)-HCl, pH 7.5, 1000 mM NaCl, 20 mM DTT, and 10 mM MgCl2 was prepared and filtered with a 0.22 μm syringe filter (Millipore Sigma) for all experiments, which was adapted from prior phase separation studies of FUS. Condensates were reconstituted by thoroughly mixing the full-length tag-free FUS at a final concentration of 10 µM or MBP FUS at a final concentration of 7.5 µM, with the phase separation buffer, an oxygen scavenging system (OSS), and a specified fluorescently labeled species (microspheres, FL mRNA, or Poly(U)_1000_ RNA). The time from assembly to data acquisition was kept under 30 minutes.

For condensate SMT experiments, samples were prepared with 10 nM AlexaFluor 488-labeled FUS and 50 pM of various fluorescent species in phase separation buffer. An OSS consisting of glucose, glucose oxidase, and catalase was employed for single-molecule tracking of RNAs. When applicable, 10% (w/v) Dextran T-500 was added as crowding agent.

### Confocal Scanning Microscopy and Circularity Analysis

Z-stack imaging (Figure 1) utilized an Alba5 confocal microscope with AlexaFluor 488 dye to visualize 3D shape and glass slide location. performed with an Alba5 time-resolved laser-scanning confocal microscope (ISS, Inc) with a pulsed supercontinuum broadband laser excitation source (Fianium WhiteLase SC-400-8-PP), avalanche photodiode (APD) detectors, and a Beckr-Hickl SPC-830 TCSPC module with a 531/40nm filter. 488 nm excitation was selected using acousto-optic tunable filters and ∼5uW average power was selected. Condensates tethered on the glass surface were found under regular scanning confocal imaging mode, and whole FOV were imaged in the x-y plane with slices taken every 20 nm starting below or at the glass surface and imaging until the focus plane was well above the condensates, ∼40 slices. These images were subjected to downstream processing and analysis. First, the raw images were deconvolved using default settings in AutoQuant software. The deconvolved images were then loading into ImageJ and thresholded before converting to binary images. The binary image was eroded to ensure edges were accurately displayed (Figure S7). The circularity of objects identified in the final images was calculated and plotted using a homemade python script.

### HILO Microscopy

Sample wells were fabricated using half-cut PCR tubes (Sigma-Aldrich) epoxy-bonded (Ellsworth Adhesives) to glass slides. For the +tether conditions only, 1 mg/mL streptavidin interface with 1 μM biotinylated FUS, anti-FUS, or DNA tethers established surface immobilization to minimize translational/rotational artifacts. After adding FUS condensates, mineral oil was layered to prevent evaporation. Imaging was performed on an Oxford Nanoimager S with a 100x 1.4 NA oil immersion super apochromatic objective, 405, 473, 532, 640 nm lasers with appropriate dual-band emission filters, and a Hamamatsu sCMOS Orca flash 4 V3 camera, using highly inclined and laminated optical sheet (HILO) microscopy [51] (52° laser angle) at 24 °C with 117 nm pixel size [52]. Intra-condensate SMT was achieved using the red (640 nm, ∼30 mW) channel. Imaging parameters were tailored to each sample type: 200 nm fluorescent beads were imaged at 100 ms exposure for main figures (Figs. 2 and 3) and at 20 ms for supplementary data; FL RNA was imaged at both 100 ms and 20 ms, and these data were further down-sampled to 200 ms for diffusion analysis (Figure 4); poly(U)_1000_ RNA was imaged at 20 ms (Figure 5); and, for the comparison of different tethering methods, FL RNA was imaged at 20 ms (Figure 5). This approach enabled direct comparison of molecular dynamics across different samples and experimental conditions while ensuring appropriate temporal resolution for diffusion analysis.

### Single-Molecule Tracking Fluctuation, Rotation, and Diffusion Analysis

Filtered videos were subjected to single-molecule tracking analysis using TrackMate, where spots were detected with a Laplacian of Gaussian (LoG) detector (object diameter: 4 pixels, 468 nm; quality threshold ≥15, determined by the peak of false-positive spot quality values). Trajectories were extracted using the Linear Assignment Problem (LAP) algorithm with a maximum linking distance of 4 pixels and no gap closing. Exported trajectories were analyzed using custom Python scripts for fluctuation, rotation, and diffusion profiling.

The *α* value was calculated by fitting the (MSD)-lag time (*τ*) curve on a log-log scale using the relation,

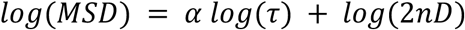

where *n* is the dimension of tracking (*n* = 2 in our case) and *D* is the diffusion coefficient. For optimal fitting [53], half the trajectory length was used unless the trajectory was shorter than 5 steps, in which case a minimum of 4 MSD-*τ* points were required. Trajectories with R² < 0.7 or with extremely small α values were excluded from further analysis.

Diffusion analysis generated distributions of mean step size, anomalous component (*α*), and apparent diffusion coefficient (*D_app_*). Mean step size was calculated for all trajectories, while α components were determined only for trajectories with a mean step size >=30nm. The *D_app_* and localization error were calculated for non-confined trajectories by fitting MSD-*τ* curve to the following formula, which is optimized for least-square fit of MSD [53,54]:

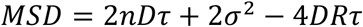

where *σ* is the localization error and *R* is the motion blur coefficient (*R* = 1⁄6 for continuous imaging).

For ensemble MSD-*τ* analysis, tracks were filtered to include only those with a minimum length of 10 frames and sufficient mobility (***log*_10_***D_app_* above the static error threshold and displacement ≥0.2 μm). Localization fluctuation analysis was performed by calculating the stepwise displacement (Δ***r***) for each trajectory, extracting the average fluctuation per track. These metrics were compared across experimental conditions using Mann-Whitney U or Kruskal-Wallis tests and Cliff’s Delta effect size.

Rotation analysis was conducted by calculating the angle between the position vector of each tracked molecule (relative to the condensate center) and its initial position vector, yielding a time series of rotation angles for each trajectory (Fig 3b). Trajectories were further analyzed to extract angular velocity 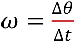 and acceleration 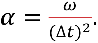

The angular velocity autocorrelation and velocity-acceleration correlation was calculated from the following equations.

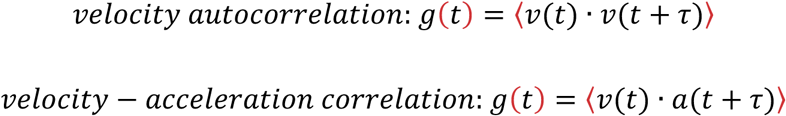

This correlation was plotted as a function of phase space for each data point in a condition to produce a useful phase space violin plot. The phase space distribution is particularly revealing because it captures the fundamental dynamical differences between tethered and untethered biomolecular condensates. This plot effectively transforms time-series rotation data into a representation that characterizes the underlying physical behavior of the system. The plot displays two key metrics for each tracked microsphere: (1) Velocity-Acceleration Correlation (x-axis) measures how angular velocity changes relate to angular acceleration. Negative values indicate damped or restrained motion (like a pendulum), while positive values suggest driven or self-reinforcing motion. (2) Phase Space Area (y-axis) quantifies the “territory” explored by the microspheres in velocity-acceleration space. Larger values indicate greater freedom of movement and more complex rotational dynamics. All analyses were implemented in homemade Python scripts.

### Monte Carlo Simulations of Single-Molecule Trajectories in tethered and untethered condensates

Our simulation framework models projected 2D single-molecule trajectories within spherical biomolecular condensates, using parameters and update logic designed to closely mimic SMT experiments. Each simulation was defined by a condensate diameter sampled from 0.3 to 5 μm and a molecular diffusion coefficient (D) sampled logarithmically from 0.001 to 10 μm²/s. The anomalous diffusion exponent (α) was set to either 1.0 (normal diffusion) or 0.5 (subdiffusive), representing key physical regimes most commonly observed in biological condensates.

To ensure efficient and uniform sampling of the (log10D, diameter) parameter space, we used Sobol low-discrepancy sampling with 200 sample points. For each parameter combination, 1,000 molecules were initialized at random positions within a square region encompassing the condensate.

Molecular motion was simulated as 2D fractional Brownian motion (FBM), generated using the Python fbm package (Davies-Harte method) [55], with the specified D and α. All simulations were performed at a time step of 50 ms for 20 steps, matching typical SMT experimental conditions.

To model the effect of thermally driven rotational dynamics of the condensate, we implemented stochastic rotation at each time step. The mathematical expectation of the angular frequency for rotational diffusion was set using the Stokes-Einstein relation for a sphere:

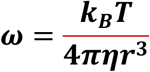

where ***k_B_*** is Boltzmann’s constant, *T* is temperature set to room temperature 298 K, ***η*** is the viscosity of the surrounding medium (fixed at 5 mPa·s, representing protein solution at around 10 mg/ml concentration), and *r* is the condensate radius in meters. At each time step, the positions of all molecules within the same condensate were rotated by a small random angle, drawn from a normal distribution with variance determined by *ω* and the time step, and scaled by the molecule’s distance from the condensate center. This rotation was applied perpendicular to the position vector, mimicking the effect of stochastic rotational diffusion of the entire droplet.

The simulations updated molecule positions at discrete time intervals (20 steps, each 50 ms), combining FBM increments and rotational displacements. This framework allows toggling rotational dynamics on or off to generate both control and experimental datasets. All simulations were performed under conditions reflecting typical SMT experiments. The resulting molecular trajectories were output for downstream analysis of apparent diffusion coefficients and anomalous exponents.

All simulation code, including parameter sweeps and Sobol sampling, was implemented in Python using numpy, fbm, and scipy libraries.

## Data & Code Availability Statement

Data are available at [to be filled]. Python scripts to process data are available at: https://github.com/walterlab-um/condensate-tether-compare-in-vitro

## Declaration of generative AI and AI-assisted technologies in the writing process

During the preparation of this work the authors used ChatGPT (OpenAI) as an editing tool to improve the readability and language of the manuscript. After using this tool, the authors reviewed and edited the content as needed and take full responsibility for the content of the published article.

## CRediT Authorship Contribution Statement

**Emily Sumrall:** Conceptualization, Methodology, Formal Analysis, Investigation, Data Curation, Writing – Original Draft, Visualization. **Guoming Gao:** Conceptualization, Methodology, Validation, Formal Analysis, Investigation, Writing – Review & Editing. **Shelby Stakenas:** Investigation, Writing – Review & Editing. **Nils Walter:** Conceptualization, Resources, Writing – Review & Editing, Supervision, Project Administration, Funding Acquisition.

## Acknowledgements

We sincerely thank Alexander Johnson-Buck, Sarah Veatch, Natalie Rogers, Adrien Chauvier, and Thanh Lai for their insightful discussions on developing the analysis pipelines for our datasets. We appreciate help from Damon Hoff at the Single Molecule Analysis in Real-Time (SMART) Center of Biophysics at the University of Michigan, for guidance on confocal scanning imaging and analysis.

## Appendix A: Supplemental Material

The following are the supplementary material to this article.

