## Supplemental Material for "Surface-Tethering Enhances Precision in Measuring Diffusion Within 3D Protein Condensates"

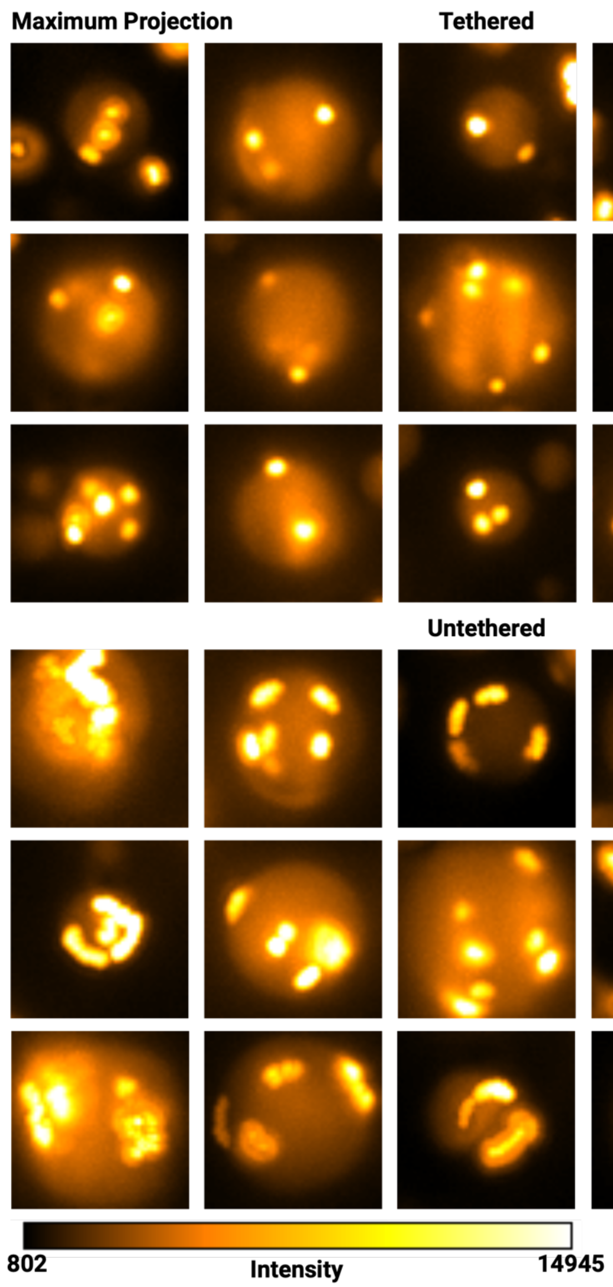


**Figure S1 | Maximum projections of fluorescent beads reveal underlying rotation and translation during imaging.** 200 nm carboxylated fluorescent beads are immobile at the 10s imaging time using 100 ms frame rate, thus maximum projections of intensity values are a simple and effective approximation for observing rotation or translation of whole condensates. **Tethered |** Maximum projections reveal stationary beads for the duration of imaging and thus indicate stably tethered condensates. **Untethered |** Maximum projections reveal diffusive beads and thus indicate unstable condensates with rotation or translation throughout the duration of imaging.

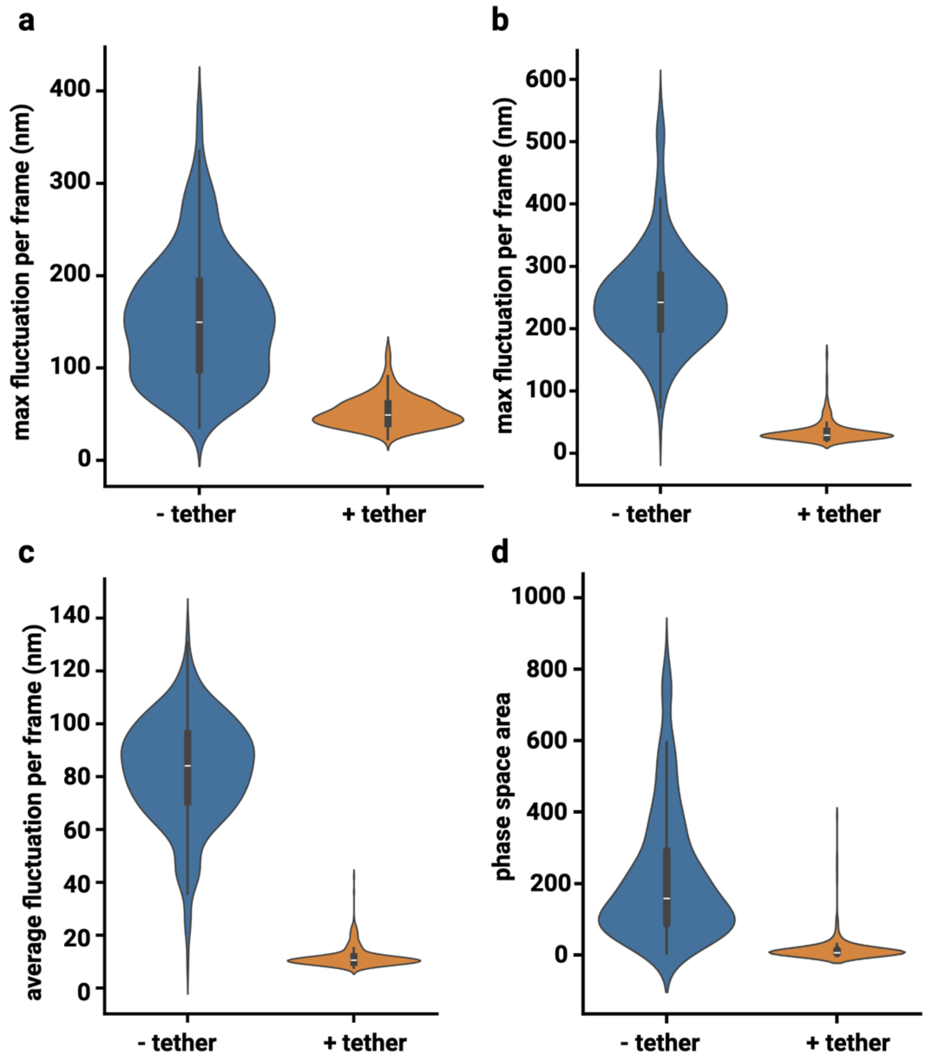


**Figure S2 | Fluctuation and rotation characteristics are independent of imaging frequency.** 200 nm carboxylated fluorescent beads imaged at 100 ms for 100 frames. **a |** Maximum frame-to-frame localization fluctuations for all condensates in each condition. **b-d |** 200 nm carboxylated fluorescent beads imaged at 20 ms for 500 frames. **b |** Maximum frame-to-frame localization fluctuations for all condensates in each condition. **c|** Average frame-to-frame localization fluctuations for all condensates in each condition. **d |** Phase space area scatter plot of velocity-acceleration correlation reveals distinct physical characteristics for untethered versus tethered condensates. As expected, these results align with Figure 3 which compare the same conditions using a 100 ms frame rate. $N_{untethered}$ = 389, $N_{tethered}$ = 310. Statistics annotation: Mann-Whitney U, ns: 0.05 < p <= 1, *: 0.01 < p <= 0.05, **: 0.001 < p <= 0.01, ***: 0.0001 < p <= 0.001, ****: p <= 0.0001.

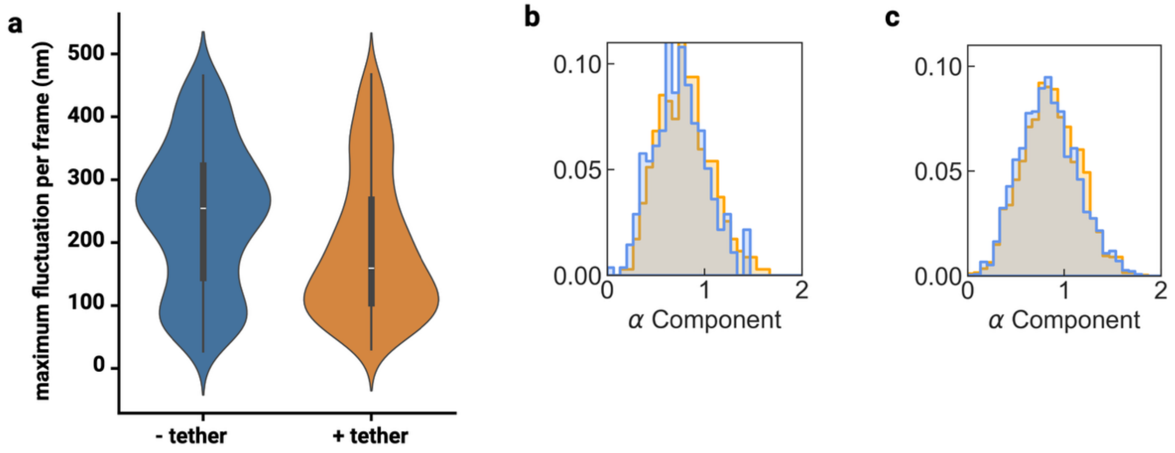


**Figure S3 | Extended SPT analysis of FL RNA in untethered or tethered condensates imaged or analyzed with 200 ms frequency. a |** Maximum frame-to-frame localization fluctuations for all condensates in each condition. **b |** Distribution of anomalous diffusion component, $\alpha$. Theoretically, an $\alpha$ value of 1 (or distribution centered around 1) reflects Brownian motion, though this value is often underestimated so a distribution around 0.7 – 1.0 is expected. **Extended SPT analysis of Poly(U)$_{1000}$ RNA in untethered or tethered condensates imaged or analyzed with 200 ms frequency. c |** Distribution of anomalous diffusion component, $\alpha$.

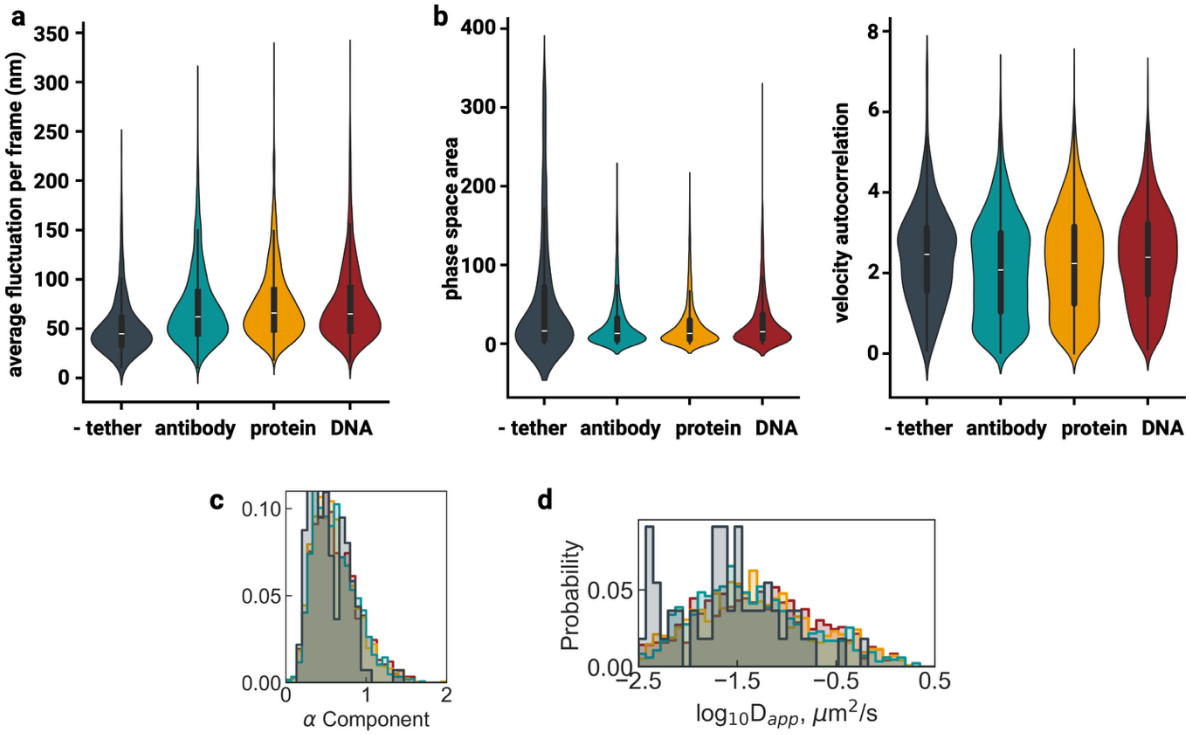


**Figure S4 | Extended SPT analysis of FL RNA in antibody-, protein-, or DNA- tethered condensates imaged or analyzed with 20 ms frequency. a |** Violin plot of average fluctuation distributions from Fig. 6a, now including the -tether condition. **b |** Violin plots of phase space area and velocity autocorrelation distributions from Fig. 6b, now including the -tether condition. **c |** Distribution of anomalous diffusion component, $\alpha$. **d |** Histogram distributions of log-transformed apparent diffusion coefficients ($\log_{10}D_{app}$, $\mu m^2/s$) from Fig. 6e, now including -tether condition.

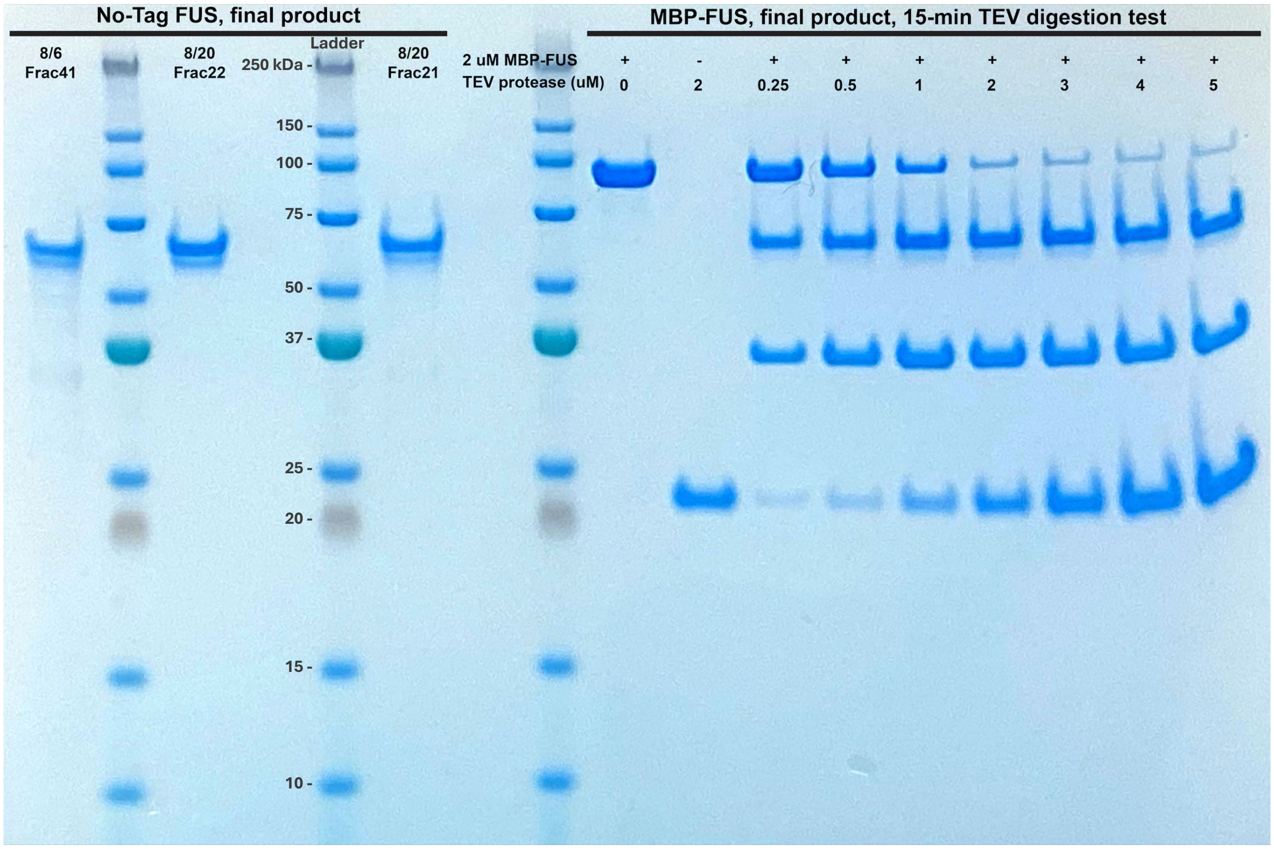


**Figure S5 | FUS purification gel. No-Tag FUS |** Occupied lanes from left to right: (1) final product fraction of no-tag FUS purification, dominant band running near ~70 kDa as expected, (2) Precision Plus Protein Ladder (Bio-rad), (3) final product fraction of no-tag FUS purification, (4) Precision Plus Protein Ladder (Bio-rad), (5) final product fraction of no-tag FUS purification. **MBP-FUS |** Occupied lanes from left to right: (1) Precision Plus Protein Ladder (Bio-rad), (2) final product fraction of MBP-FUS purification, 0 μM TEV-protease, dominant band running near ~98 kDa as expected, (3) 0 μM MBP-FUS, 2 μM TEV-protease, dominant band running near ~23 kDa as expected, (4-10) 2 μM MBP-FUS with increasing concentrations of 0.25, 0.5, 1, 2, 3, 4, 5 μM TEV-protease respectively, dominant bands running near ~98 kDa for uncleaved MBP-FUS, ~70 kDa for cleaved full length FUS, ~35 kDa for cleaved MBP fragment, and ~23 kDa as expected for TEV protease. For all lanes, signal is from Coomassie Blue staining.

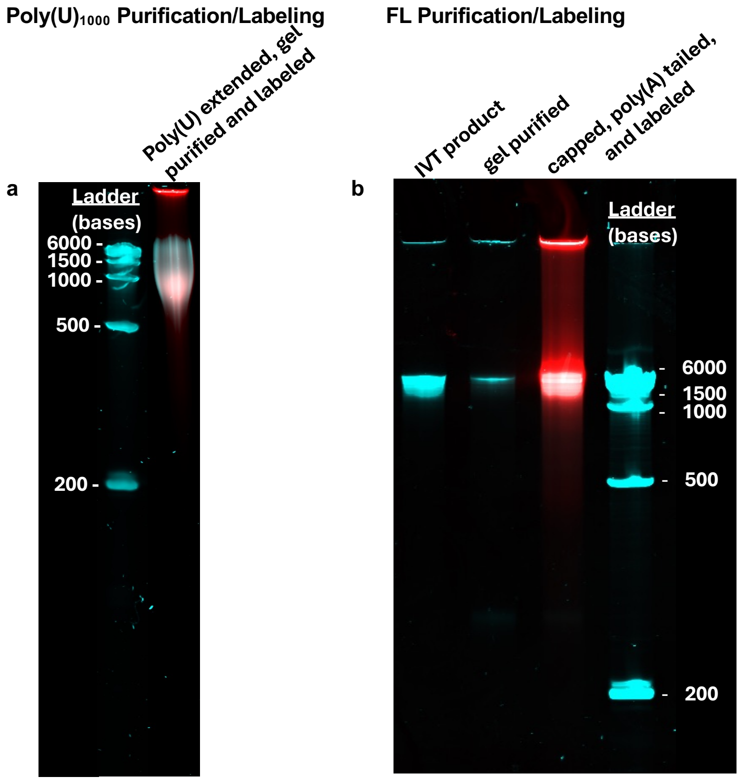


**Figure S6 | Poly(U)$_{1000}$ and FL RNA purification Gel. a |** Occupied lanes from left to right: (1) RiboRuler High Range (ThermoFisher) ladder for RNA PAGE. (2) Poly(U)$_{1000}$ gel purification product after Poly(U) extension, Azido-ATP addition, and AlexFluor-647 Click-chemistry labeling. **b |** Occupied lanes from left to right: (1) RNA IVT product (2) RNA gel purification product after IVT (3) RNA gel purification product after Azido-ATP addition, Poly(A) tail extension, 5'-OMe cap, and AlexFluor-647 Click-chemistry labeling. (4) RiboRuler High Range (ThermoFisher) ladder for RNA PAGE. For all lanes, cyan represents syber-gold stain signal and red corresponds to AlexaFluor-647 signal.

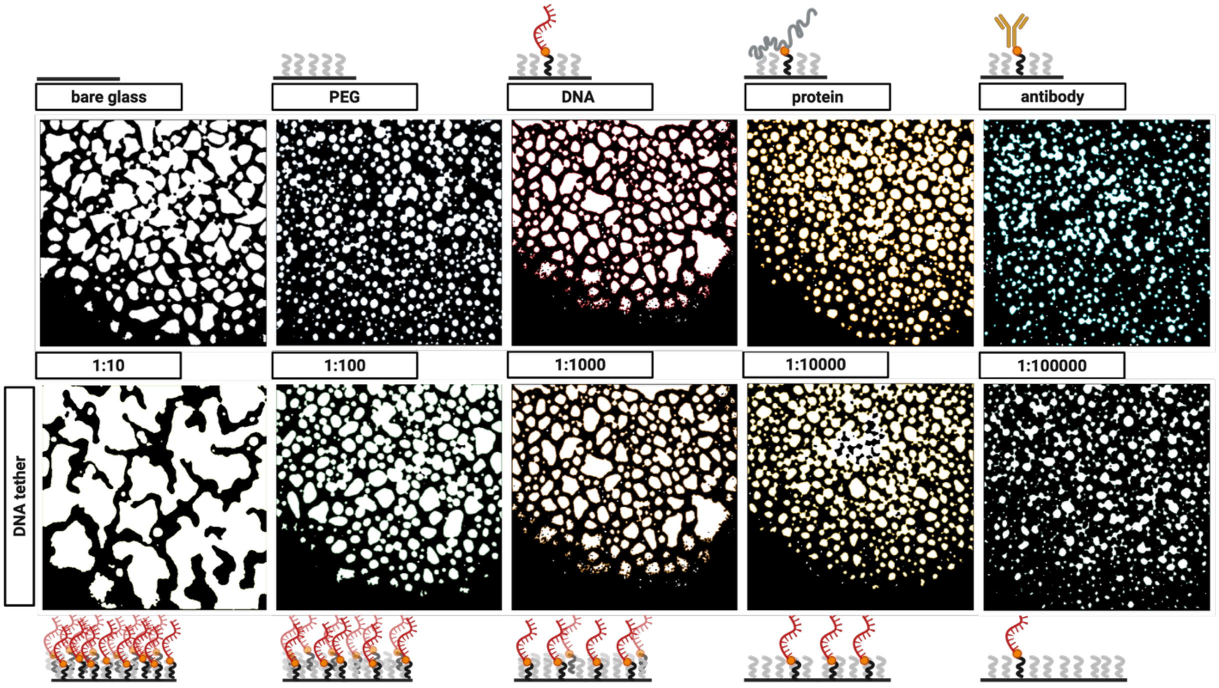


**Figure S7 | Binary threshold images with drawn contours for circularity analysis.** Full field of view is shown here and used for analysis in Fig. 1 b and d.
